# Vesicular glutamate release is necessary for neural tube formation

**DOI:** 10.1101/2025.01.05.631426

**Authors:** Raman Goyal, Patricio A. Castro, Jacqueline B. Levin, Sangwoo Shim, Grace Or Mizuno, Lin Tian, Laura N. Borodinsky

## Abstract

The brain and spinal cord originate from a neural tube that is preceded by a flat structure known as the neural plate during early embryogenesis. In humans, failure of the neural plate to convert into a tube by the fourth week of pregnancy leads to neural tube defects (NTDs), birth defects with serious neurological consequences. The signaling mechanisms governing the process of neural tube morphogenesis are unclear. Here we show that in *Xenopus laevis* embryos, neural plate cells release glutamate during neural plate folding in a Ca^2+^ and vesicular glutamate transporter-1 (VGluT1)-dependent manner. Vesicular release of glutamate elicits Ca^2+^ transients in neural plate cells that correlate with activation of Erk1/2. Knocking down or out VGluT1 leads to NTDs through increased expression of Sox2, neural stem cell transcription factor, and neural plate cell proliferation. Exposure during early pregnancy to neuroactive drugs that disrupt these signaling mechanisms might increase the risk of NTDs in offspring.

## Introduction

Nervous system development progresses through several milestones that rely on mechanisms that are believed to be gradually acquired by neural cells as they differentiate and become more specialized in their function. One of these milestones is the formation of the neural tube, which constitutes the presumptive brain and spinal cord, that originates from a flat layer of neuroectodermal stem cells called the neural plate. This process is complex and its failure leads to birth defects known as neural tube defects (NTDs) that can be lethal or result in various neurological disabilities ^1,2^, underscoring the importance of understanding the mechanisms governing neural tube formation.

The cellular events that are required for the morphogenesis of the neural tube include apical constriction of neural plate cells to enable bending, cell elongation to facilitate the curvature needed for the reshaping of the tissue and collective migration of cells towards the midline for fusion and closure of the neural tube ^2–8^. This is only an incomplete list of the multitude of cellular events that are required for neural tube formation, which points to the requirement for spatiotemporal coordination among these processes. Although several molecules that participate in each of these cellular events have been identified, the understanding of the signaling mechanisms that operate in neural plate cells is still incomplete.

In addition, during neural tube morphogenesis, an optimal number of neural plate cells needs to be available. Indeed, the role of cell proliferation in neural tube formation has been a subject of extensive investigation ^9–18^. These studies present evidence that cell cycle perturbation at specific developmental stages to either increase or decrease neuroepithelial cell proliferation can lead to neural tube defects in various species. Interestingly, in *Xenopus laevis* embryos, blocking cell proliferation by inhibiting DNA synthesis following gastrulation does not interfere with neural tube closure ^11,18^. In contrast, inhibiting cell cycle progression in embryos in which genetic (curly tail mouse; ^19^) or pharmacological (valproic acid- incubated *Xenopus laevis* embryo; ^20^) perturbations induced imbalanced neural plate cell proliferation, partially rescues the NTD phenotype. Overall, these studies suggest that precise regulation of neural plate cell proliferation is important for the morphogenesis of the neural tube.

The causes of NTDs appear to be both genetic and environmental. The challenges in identifying the etiology of these birth defects are likely related to the diversity of cellular processes involved which may be regulated by different mechanisms. One of the environmental factors that is associated with increased risk of NTDs is *in utero* exposure of the fetus to antiepileptic drugs during early pregnancy ^21–26^. Although the prevailing view has been that these drugs interfere with the morphogenesis of the neural tube through off-target effects ^27^, our previous study has demonstrated that valproic acid, a commonly used antiepileptic drug, induces NTDs in frog embryos by a similar cellular mechanism as interfering with NMDA receptor-dependent glutamate signaling, which perturbs regulation of neural plate cell proliferation. Moreover, valproic acid-induced NTDs are partially rescued not just by inhibiting cell proliferation, but also by enhancing glutamate signaling ^20^. These studies suggest that the regulation of cell proliferation that is important for neural tube folding is dependent on glutamate signaling.

Our previous study has shown that indeed, NMDA receptors are expressed and their inhibition or knockdown results in a decrease in Ca^2+^ transients present in neural plate cells during tissue folding and in NTDs ^20^. However, the mechanism by which glutamate is released in a developmental stage when neurons are not yet fully differentiated, and synapses are not formed is unclear. Synaptic glutamate release is dependent on the expression and compartmentalization of key molecules that participate in the storing of glutamate in synaptic vesicles, anchoring of vesicles in synaptic terminals, fusion of vesicles and release of neurotransmitter ^28^. Prominently, vesicular glutamate transporters are indispensable for filing synaptic vesicles with glutamate in glutamatergic synapses ^29^. Whether aspects of the mechanism of vesicular glutamate release in synapses are also recruited at the early stages of neural plate folding has never been investigated before.

Here we show that the vesicular glutamate transporter 1 (VGluT1) is expressed in the neural plate and is necessary for neural tube formation. This study demonstrates VGluT1-dependent release of glutamate that mediates Ca^2+^ dynamics to regulate neural plate cell proliferation and morphogenesis of the neural tube.

## Results

### VGluT1 is expressed in the folding neural plate and mediates glutamate release

To investigate the mechanisms of glutamate release in the neural plate we examined expression of VGluT1, the vesicular glutamate transporter known to be expressed in *Xenopus laevis* ^30–32^. mRNA expression and Western blot assays show that VGluT1 is expressed in neural plate stages (Fig. 1A-B) , in agreement with previous studies ^31^, while transcript for the excitatory amino acid transporter 5 (*eaat5*) known to be aa retina-specific glutamate transporter ^33^, is not detected during neural plate stages (Fig. 1A). VGluT1 protein localizes to neural plate cells and the number of VGluT1+/Sox2+ neural plate cells increases as neural plate folding progresses (Fig. 1C). The decreased protein level when knocking down VGluT1 expression proves the specificity of immunodetection (Fig. 1B,D). In addition, transcripts for other proteins associated with neurotransmitter vesicular release, such as syntaxin 1A (*stx1a*), vesicle associated membrane protein 1 (*vamp1*), and synaptosome associated protein 25 (*snap25*), are also detected at these neural plate stages (Fig. 1A), suggesting that the molecular machinery for vesicular glutamate release is available during neural tube formation. Indeed, ultrastructural assays we performed at these stages show that vesicular structures are prominent in neural plate cells (Fig. 1E-E’) and that VGluT1 colocalizes in subcellular regions of potential vesicular glutamate release (Fig. 1F-F’).

**Figure 1.**
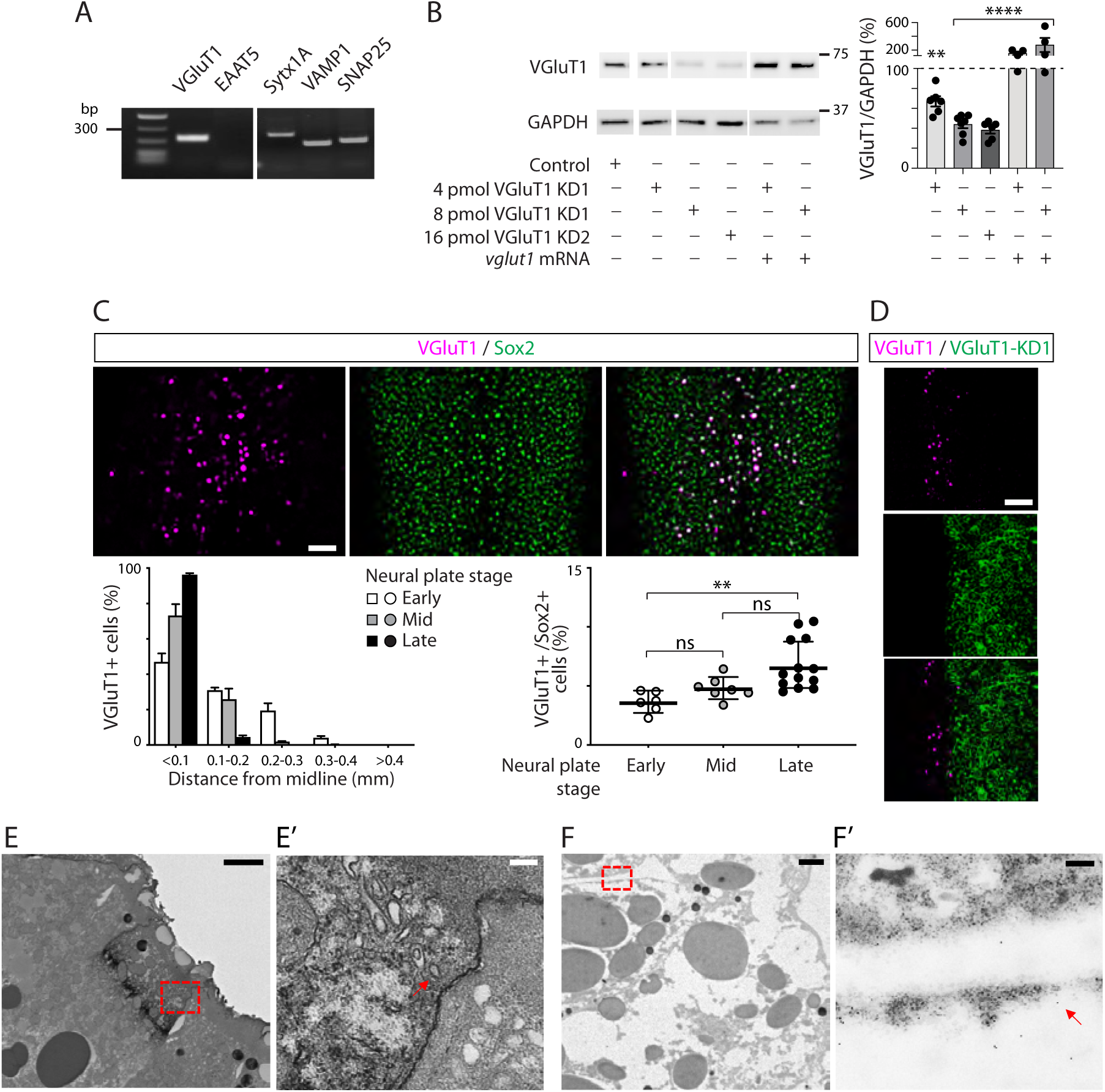
VGluT1 is expressed in the neural plate of *Xenopus laevis* embryos. **A.** RNA was isolated from embryos at mid-neural plate stages and reverse transcriptase-PCR assays were performed for *vglut1*, *eeat5*, *syntaxin1a* (*stx1a*), *vamp1* and *snap25*. **B.** Western blot assays were performed in whole-cell lysates from neural-plate-stage control, VGluT1-knockdown (KD) and *vglut1* mRNA-injected embryos. GAPDH was used as loading normalizer. Shown is a representative example. Graph shows normalized signal intensities from individual samples and means as percentages of control values (dashed line) from N≥4 experiments. Statistical analysis was done with one-sample t and Wilcoxon test, **p<0.01, ****p<0.0001. **C.** Representative maximum intensity projection of whole-mount immunostained dorsal half of mid-neural plate stage embryo for VGluT1 and Sox2 (neural stem cell marker). Graphs show mediolateral (left; mean percent of VGluT1+ cells per 100-μm bin from the midline compared to the total number of VGluT1+ cells ± SEM) or developmental (right; individual data and mean percent of VGluT1+ cells of the total number of Sox2+ cells ± SEM) distribution of number of VGluT1 immunopositive (+) cells. **p<0.01, ns: not significant, one-way ANOVA. N=3. **D.** VGluT1-KD decreases the number of VGluT1+ cells in the neural plate. Shown is a maximum intensity projection of whole-mount immunostained unilaterally VGluT1-KD neural plate from a representative embryo, N=3. **E**-**F**. Neural plate stage embryos were processed for transmitted (**E**) and immuno-transmitted (**F**) electron microscopy assays. Shown are representative examples. Dashed boxes in (**E**,**F**) indicate fields of view shown in (**E’**,**F’**), respectively. Arrow in (**E’**) points to vesicular structures and in (**F’**) to structure with lipidic-looking background (vesicular) close to the cell-cell border immunopositive for VGluT1, N=3. Scale bars, 2 (**E**,**F**) and 0.2 (**E’**,**F’**) μm.

To determine whether expression of VGluT1 is functional and necessary for glutamate release, we performed live imaging of intact *Xenopus laevis* embryos expressing a genetically encoded sensor of released glutamate, iGluSnFR (123D5-spacer-modified; Lin Tian’s lab). We confirmed that the sensor reports levels of extracellular glutamate concentration by adding glutamate to the bathing solution, which results in an increase in iGluSnFR signal (Fig. 2A). Results show that the sensor reports a stronger signal in the neuroectoderm compared to the non-neural ectoderm, suggesting higher glutamate release in the neural plate (Fig. 2B). In contrast, VGluT1 knockdown decreases iGluSnFR signal but not membrane-mCherry fluorescence in the affected neural plate (Fig. 2C). This decrease is not due to reduced expression of iGluSnFR because staining for level of expressed reporter (myc-tagged) reveals that expression is comparable in wild type and knockdown halves of neural plate (Fig. 2D). These results indicate that VGluT1 is expressed and necessary for glutamate release from neural plate cells.

**Figure 2.**
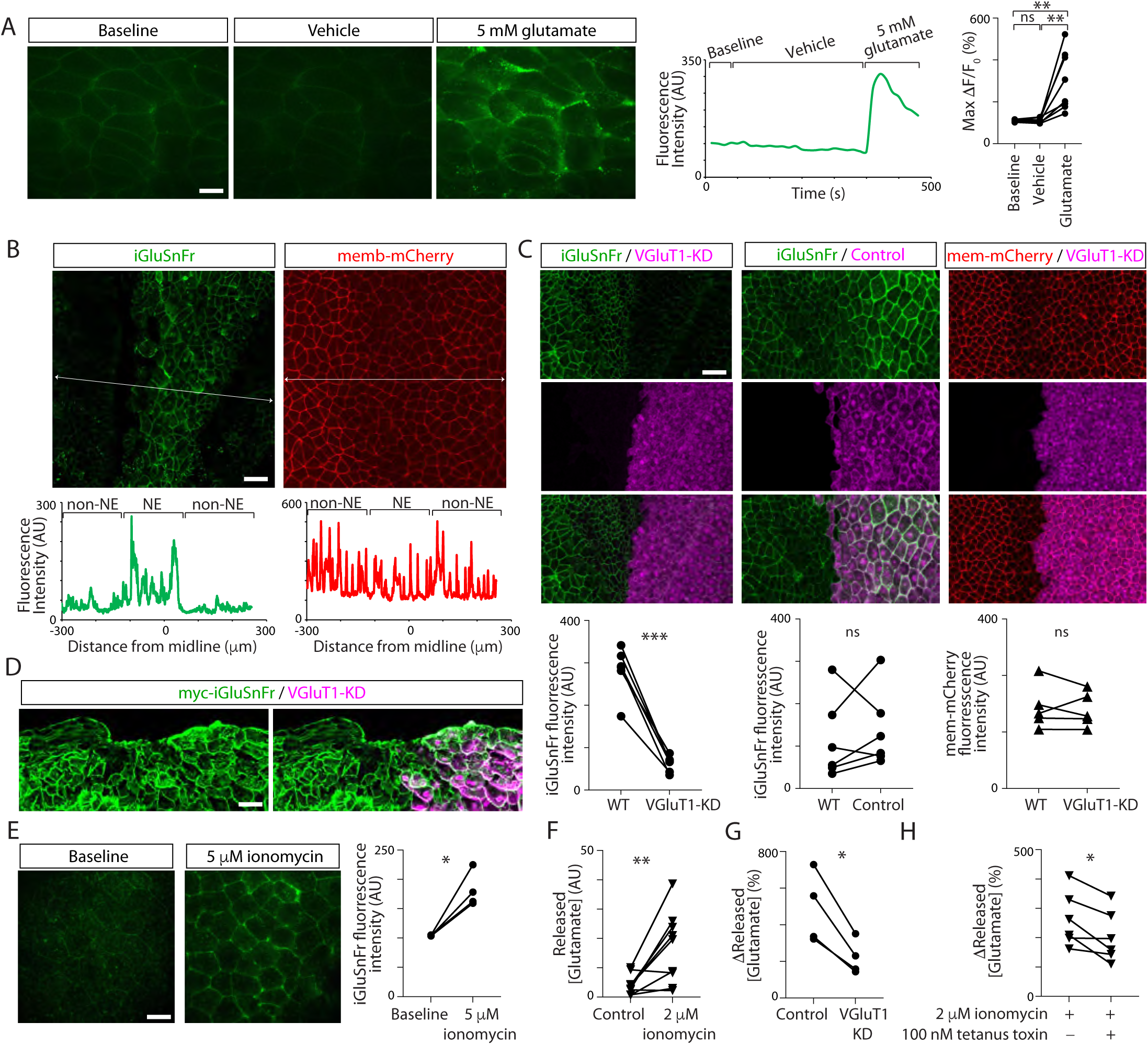
**Glutamate is released from neural plate cells in a VGluT1- and Ca^2+^-dependent manner**. **A**-**E**. Two-cell stage embryos were bilaterally microinjected with mRNA encoding iGluSnFR (**A**-**E**) or membrane (memb)-mCherry (**B**-**C**) and unilaterally injected with VGluT1-morpholino 1 (VGluT1-knockdown (KD), **C**-**D**) or control-morpholino (Control, **C**). Embryos were live imaged (**A**-**C**, **E**) or processed for immunostaining (**D**). **A.** iGluSnFR expressed in the neural plate senses extracellular glutamate levels. Shown are single time frames of time-lapse recording of the neural plate in whole embryo when either vehicle or 5 mM glutamate was added to the bathing solution. Trace shows representative change in iGluSnFR fluorescence intensity over time. Graph shows individual maximal change in iGluSnFR fluorescence intensity after addition of vehicle or 5 mM glutamate. N=7, **p<0.01, ns: not significant, paired ANOVA. **B.** Glutamate is released from neural plate cells. Shown is maximum intensity projection of confocal image of iGluSnFR or memb-mCherry-expressing embryos and mediolateral fluorescence intensity profile for both reporters. While iGluSnFR exhibits higher intensity in membrane of neural plate cells located within 200 μm of the midline compared to non-neural ectodermal cells, memb-mCherry does not show a differential mediolateral distribution. N=3. **C.** VGluT1-KD impairs glutamate release. Images are representative examples of maximum intensity projection of unilaterally manipulated embryos as indicated. Graphs represent iGluSnFR (left and middle) or memb-mCherry (right) fluorescence intensity of individual embryos in WT and manipulated halves of neural plate. ***p<0.001, ns: not significant, N=5 (VGluT1-KD iGluSnFR and memb-mCherry), N=6 (Control), 2-way paired t-test. **D.** Expression of iGluSnFR is not affected by VGluT1-KD. Shown is a representative example of 10 μm section of unilaterally VGluT1-KD neural plate immunostained for myc-tag linked to iGluSnFR construct, N=3. **E.** Ca^2+^ entry in neural plate cells increases glutamate release. Images show iGluSnFR expressing neural plate before and after addition of 5 μM ionomycin. Graph shows individual data of % maximal change in iGluSnFR fluorescence intensity after addition of ionomycin. N=4, *p<0.05, one-sample t and Wilcoxon test, compared to the hypothetical value of 100 (before addition of ionomycin). **F**-**H**. Neural plate stage wildtype (**F**,**H**) and bilaterally morpholino- injected embryos (**G**, Control and VGluT1-KD) were incubated with 2 μM ionomycin (**F**-**H**) or vehicle (**F**) in the presence (**H**) or absence (**F**-**H**) of 100 nM tetanus toxin for 20 min. Bathing solution was collected and released glutamate measured with a Fluorometric Glutamate Assay Kit. **F.** Graph shows released glutamate concentration in Control (vehicle) and ionomycin treated embryos, N=9, **p<0.01, 2-way paired t-test. **G.** Graph shows individual % change in released glutamate concentration in Control and VGluT1-KD embryos treated with ionomycin and compared to vehicle-treated (100%), N=4, *p<0.05, 2-way paired t-test. **H.** Graph shows individual % change in released glutamate concentration in embryos treated with ionomycin in the absence or presence of 100 nM tetanus toxin compared to untreated (100%), N=6, *p<0.05, 2-way paired t-test.

To further confirm release of glutamate from the neural plate and determine whether it is dependent on Ca^2+^ influx, as in synapses, we measured in whole live embryos glutamate release in the presence or absence of ionomycin, a Ca^2+^ ionophore, using iGluSnFR (Fig. 2E) or a fluorometric ELISA assay (Fig. 2F). Results show that Ca^2+^ influx increases glutamate release from neural plate stage embryos (Fig. 2E-F), while knocking down VGluT1 expression (Fig. 2G) or blocking vesicular neurotransmitter release by pre- incubating neural plate stage embryos with tetanus toxin (Fig. 2H) inhibits ionomycin-induced glutamate release (Fig. 2G-H). This indicates that mechanisms of Ca^2+^-mediated vesicular glutamate release are present in neural plate stage embryos.

### VGluT1 is necessary for Ca^2+^ dynamics and signaling in neural plate cells

Neural plate cells exhibit Ca^2+^ transients ^20,34,35^ that increase in frequency with the progression of neural plate folding ^20^, and are important for the morphogenesis of the neural tube ^34,35^. The increase in Ca^2+^ transient frequency in the neural plate correlates with progressive increase in the number of VGluT1- expressing neural plate cells (Fig. 1C). Hence, we assessed whether VGluT1 regulates neural plate cell Ca^2+^ dynamics. We find that knocking down VGluT1 expression in half of the neural plate almost completely abolishes Ca^2+^ transients compared to the counterpart wild type neural plate (Fig. 3A), demonstrating a dependence on VGluT1-enabled glutamate signaling for Ca^2+^ dynamics in the neural plate.

**Figure 3.**
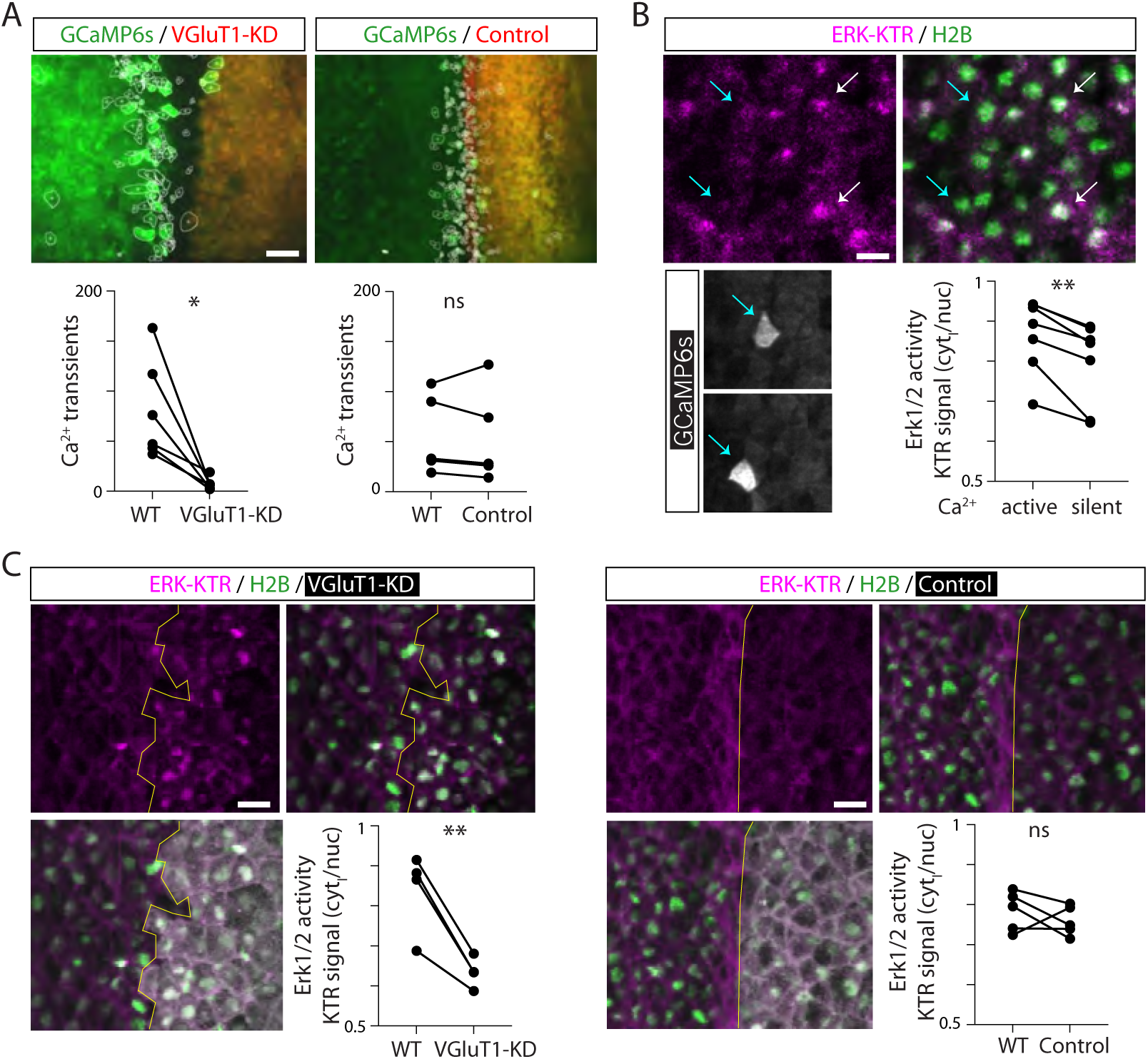
Ca^2+^ dynamics in the neural plate during folding depend on VGluT1 and activate Erk1/2. **A.** Two- cell-stage embryos were unilaterally VGluT1 knocked down (KD) and bilaterally injected with mRNA encoding GCaMP6s, Ca^2+^ reporter. Neural plate stage embryos were timelapse imaged for 5 min and number of Ca^2+^ transients were measured. Image shows representative example of unilateral (red) VGluT1- KD and Control embryos. Circled cells are those exhibiting Ca^2+^ transients during recording. Graphs show individual number of Ca^2+^ transients per 5 min in WT and VGluT1-KD or Control halves of the neural plate. N=6 VGluT1-KD and 5 Control, *p<0.05, ns: not significant, 2-way paired t-test. **B.** Two-cell stage embryos were bilaterally injected with ERK-KTR-iRFP670 (Erk1/2 activity reporter), H2B-RFP (nuclear marker), and GCaMP6s. Neural-plate-stage embryos were time-lapse imaged for 2 h, and cells exhibiting Ca^2+^ transients and those matching contralaterally their mediolateral and anteroposterior locations were selected for measurement of the ERK-KTR nuclear (delimited by H2B signal) and non-nuclear (cytosolic, delineated by GCaMP6 signal) fluorescence signal. Upper images are a representative example of single time frame of an embryo live-imaged. Cyan arrows point to two cells exhibiting Ca^2+^ transients (active), and white arrows point to similarly positioned (mediolaterally and anteroposteriorly) cells that did not exhibit Ca^2+^ transients (silent) during recording. Bottom images show the two cells exhibiting Ca^2+^ transients during peak of transient. Graph shows mean ratio of ERK-KTR cytosolic/nuclear fluorescence intensity per embryo in active and silent cells. N=7 embryos, n=10 cells per embryo, **p<0.01, 2-way paired t-test. **C.** Two-cell stage embryos were bilaterally injected with ERK-KTR-iRFP670 and H2B-RFP, and unilaterally with VGluT1- KD or Control along with tracer. Neural plate stage embryos were time-lapse imaged and ERK-KTR nuclear (delimited by H2B signal) and cytosolic fluorescence signal was measured in WT and injected neural plate cells. Images show single time frame of unilateral VGluT1-KD and Control embryos. Yellow line separates WT from injected half neural plate. Graph shows mean ratio of ERK-KTR cytosolic/nuclear fluorescence intensity in each half of the neural plate per embryo. N=4 embryos, n=10 cells per embryo (VGluT1-KD) and N=5 embryos, n=10 cells per embryo (Control), **p<0.01, ns: not significant, 2-way paired t-test.

Glutamate-mediated signaling is known to recruit various Ca^2+^-regulated pathways, including the MAPK cascade ^36,37^. Our previous study demonstrated that NMDA receptor activation elicits Erk1/2 phosphorylation in neural plate cells ^20^. We examined the status of Erk1/2 activity in neural plate cells by simultaneously live imaging Erk1/2 and Ca^2+^ activity in intact, neurulating embryos expressing specific, genetically-encoded reporters. We first determined that the Erk1/2 reporter (ERK-KTR ^38^) expressed in *Xenopus laevis* embryonic spinal cord and muscle cells reports on changes in Erk1/2 activity elicited by an agonist or antagonist of the pathway (Extended Data Fig. 1). We find that cells with Ca^2+^ transients (active) exhibit higher levels of Erk1/2 activity than those cells without Ca^2+^ transients (silent) during the recording (Fig. 3B). Importantly, VGluT1-deficient neural plate cells exhibit decreased Erk1/2 activity compared with the wild type counterparts (Fig. 3C), suggesting that VGluT1-mediated glutamate release in the neural plate activates Erk1/2.

### VGluT1 is necessary for neural tube formation through the regulation of neural plate cell proliferation

The data thus far demonstrates that VGluT1 mediates glutamate release in the folding neural plate that is necessary for recruiting Ca^2+^ signaling during neural tube formation. We therefore investigated the importance of VGluT1-dependent glutamate release in neural tube closure. By knocking down (Fig. 1B) or knocking out (via CRISPR/Cas9) VGluT1 expression (Extended Data Fig. 2) in developing embryos we find that VGluT1 deficiency leads to NTDs in *Xenopus laevis* embryos that manifest as failure of closure of the neural tube in the midline (Fig. 4). The penetrance of VGluT1 knockdown-induced NTD phenotype correlates with the extent of knockdown (Fig. 1B) and is rescued by restoring VGluT1 expression (Fig. 4), indicating specificity of VGluT1-KD-elicited NTD phenotype.

**Figure 4.**
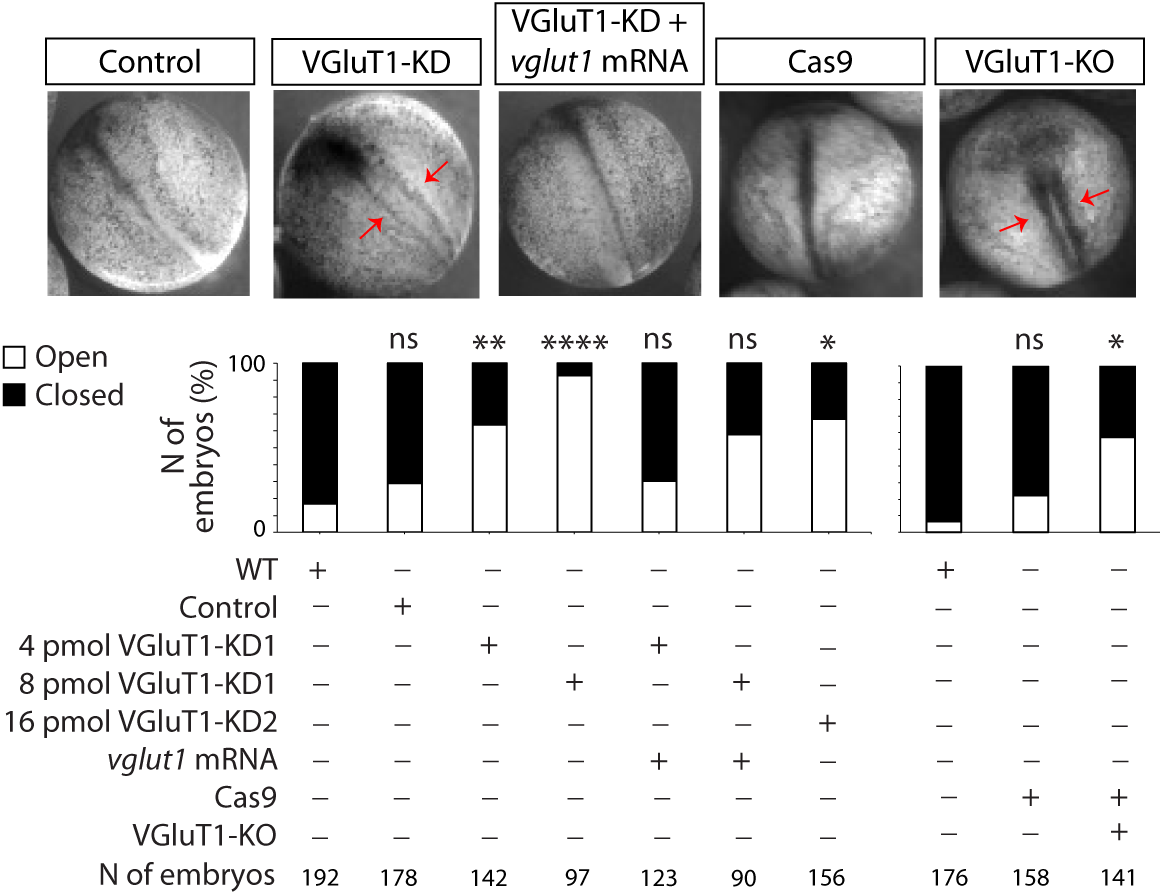
VGluT1 is necessary for neural tube formation. Two-cell stage embryos were bilaterally injected with 2-4 pmol VGluT1-morpholino 1 translation-blocking (VGluT1-KD1), 8 pmol VGluT1-morpholino 2 splicing-blocking (VGluT1-KD2) or 8 pmol control-morpholino (Control) without or with morpholino- resistant *vglut1* mRNA, or with Cas9 or Cas9 + VGluT1 sgRNA (VGluT1-KO). Images show representative examples of embryos at the time neural tube closed in Control group. Red arrows indicate open neural tube. Graph shows % of embryos with open and closed neural tubes in each group. N≥5 experiments, n of embryos indicated in graph for each group, *p<0.05, **p<0.01, ****p<0.0001, ns: not significant, one-way ANOVA, mixed-effects analysis with Geisser-Greenhouse correction, Dunnett’s multiple comparisons test, compared to WT.

We find that most VGluT1-expressing cells are non-proliferative as revealed by the small proportion of VGluT1/PCNA or VGluT1/PHH3 co-immunopositive neural plate cells during neural plate folding (Fig. 5A). Hence, we examined whether VGluT1 is necessary for regulating neural plate cell proliferation during neural tube formation. Results show that VGluT1 knockdown increases the number of proliferative cells in the neural plate, evidenced by higher number of total Sox2- and Sox2/PCNA-expressing cells (Fig. 5B). Moreover, Western blot assays reveal that VGluT1 knockdown increases Sox2 expression levels (Fig. 5C). Altogether, these results indicate that VGluT1 regulates neural plate cell proliferation to enable the morphogenesis of the neural tube.

**Figure 5.**
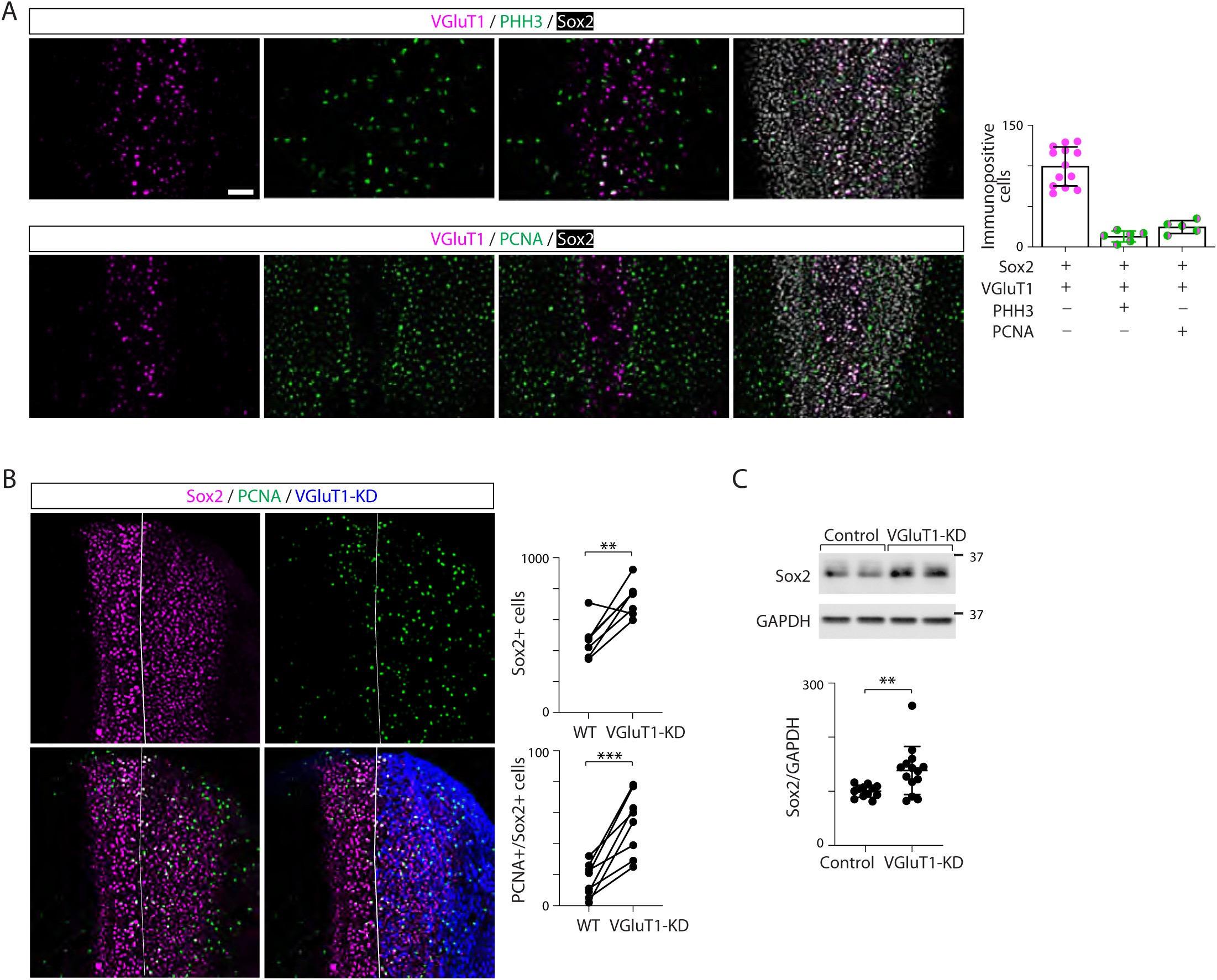
VGluT1 regulates neural plate cell proliferation and Sox2 expression. **A.** Wild type neural plate stage embryos were processed for whole-mount immunostaining for VGluT1, Sox2 and cell proliferation markers PCNA and PHH3. Shown are representative images. Graph shows individual and mean±SD number of total and proliferation marker immunopositive VGluT1 neural plate cells (Sox2+). **B.** Two-cell stage embryos were unilaterally injected with VGluT1-KD along with tracer. Neural plate stage embryos were processed for immunostaining. Images show representative example. Graphs show individual number of immunopositive (+) cells in both halves of the neural plate per embryo. N=7 embryos, **p<0.01, ***p<0.001, 2-way paired t-test. **C.** Two-cell stage embryos were bilaterally injected with VGluT1-KD or Control. Neural plate stage embryos were processed for Western blot assays for Sox2 and GAPDH, as loading control. Image is a representative example. Graph shows individual samples and mean±SD normalized Sox2 levels. N=5 experiments, **p<0.01, 2-way t-test.

## Discussion

The vesicular release of neurotransmitter has been primarily studied in the context of chemical synapses between neurons and target cells. Nevertheless, neurotransmitters and their synthetic and release machineries are known to be expressed prior to synapse formation ^31,39,40^. It could be argued that this precedent is needed for preassembling the specialized molecular platform required in synapses. Alternatively, it could indicate that neurotransmitter signaling plays a critical role even at early stages of nervous system development like the neural plate period. This study demonstrates that the vesicular release of glutamate in the neural plate occurs through mechanisms in part shared by those in synapses, and that this release is indispensable for the formation of the neural tube. These findings attest that expression of glutamate release machinery is not merely preparatory for the synaptogenic period occurring later in development, but that the signaling it elicits is essential for regulating neural cell proliferation during neural plate folding.

Here we show expression of components of the vesicular, Ca^2+^-regulated synaptosomal associated protein receptor (SNARE) complex machinery and VGluT1 immunogold particles localizing to vesicular ultrastructure in the proximity of the apicolateral membrane of neural plate cells. Moreover, we show that glutamate release from the neural plate is VGluT1-dependent and enhanced by Ca^2+^ influx. This suggests that the SNARE-dependent vesicular mechanism is present in neural plate cells. The extent to which this mechanism of vesicular glutamate release in neural plate cells resembles the quantal release seen in synapses requires further investigation. Nevertheless, it appears that expression of vesicular release machinery and ultrastructure characteristic of dynamic exocytic and endocytic events, indicative of vesicular release, are staples of the folding neural plate across vertebrates ^3,7,8,41–44^. The morphogenesis of other tissues is also dependent on exocytosis, including the dorsal mesoderm in *Xenopus laevis* ^45^ and the *Drosophila* air sac primordium ^46^, where vesicular release of glutamate through SNARE-dependent contacts between cytonemes protruding from disc cells and air sac primordium cells has been reported ^46^. Therefore, specialized vesicular glutamate release might be a key mechanism that enables embryonic morphogenesis.

Vesicular release is a more energetically demanding process than alternative, non-quantal release mechanisms, but offers more spatiotemporal regulation and organization of the signaling. The presence of this mechanism in the neural plate suggests that this precision in signaling enabled by the vesicular release of glutamate is important for the specific cellular processes taking place during neural plate folding. Indeed, we find that Ca^2+^ transients, which are strongly dependent on VGluT1-mediated glutamate release, initially occur in individual neural plate cells, with clusters of coactive cells only becoming apparent later in the folding process ^20,34,35^. This suggests that the signaling is initially targeted to individual cells during the onset of neural plate folding, allowing for cell autonomy by restricting the release of glutamate to specific cells and subcellular domains.

The locally released glutamate and subsequent Ca^2+^ dynamics in individual cells may be necessary for cell- autonomously regulating the cellular behaviors necessary for their participation in the folding of the neural plate. These include decisions on whether a cell should exit the cell cycle to change shape and migrate or continue proliferating to achieve the necessary cell count for neural tube morphogenesis. The VGluT1- dependent Ca^2+^-Erk1/2 activity that neural plate cells exhibit, demonstrated here, together with previous findings showing that the NTD phenotype from NMDA receptor knockdown is rescued by expressing an inducible, constitutively active MAPK during neural tube formation ^20^, suggest that Erk1/2 is downstream glutamate-dependent Ca^2+^ signaling in the regulation of neural plate cell proliferation necessary for neural tube morphogenesis (Fig. 6).

**Figure 6.**
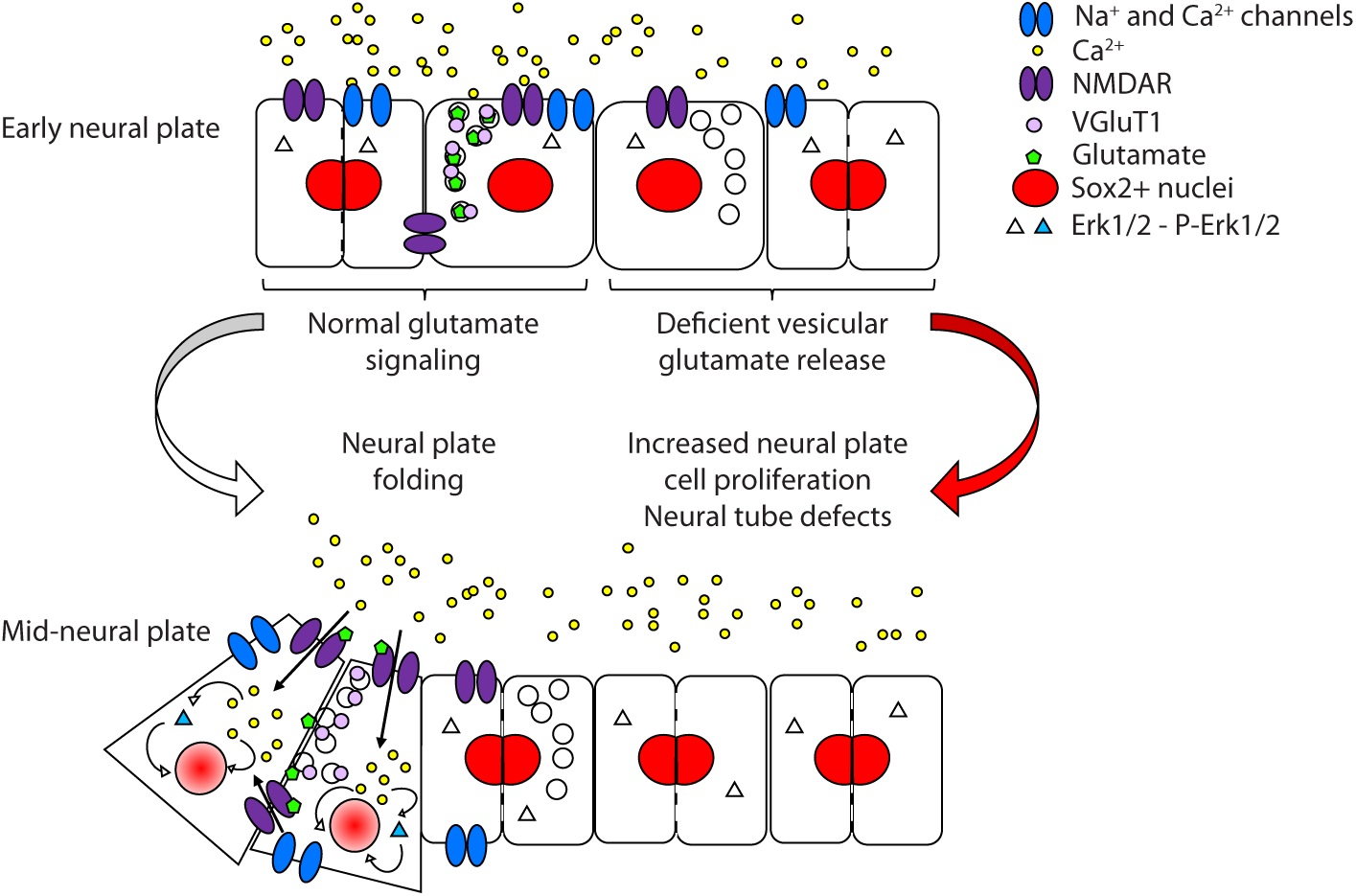
Model of mechanism of VGluT1-dependent regulation of neural tube formation. Vesicular release of glutamate from neural plate cells is necessary for the regulation of neural plate cell proliferation through the recruitment of Ca^2+^ and Erk1/2 signaling and downregulation of Sox2 expression. This in turn allows for the timely folding of the neural plate during neural tube morphogenesis.

Sox2 is a crucial transcription factor for the maintenance of neural cell stemness ^47^ by supporting neural stem cell self-renewal in the developing and adult nervous system ^48,49^. Its expression is tightly regulated and alterations in this regulation has profound consequences, with constitutive expression inhibiting neuronal differentiation and downregulation leading to premature cell cycle exit and neurogenesis ^48^. We demonstrate that proper Sox2 levels in neural plate cells are dependent on VGluT1 expression which indicates that released glutamate regulates Sox2 expression. This finding reveals a mechanism by which glutamate signaling regulates neural plate cell proliferation. Given that Sox2 expression is negatively correlated to the frequency of Ca^2+^ transients during neural plate folding ^20^, it suggests that Ca^2+^ signaling could regulate Sox2 levels transcriptionally, as previously shown for Ca^2+^-dependent regulation of Sox2 expression in *Xenopus* developing neural tube ^50^, and/or post-translationally, through phosphorylation- dependent mechanisms ^51^. We propose a model where VGluT1-dependent, Ca^2+^ mediated release of glutamate in selected neural plate cells signals the need for cells to withdraw from the cell cycle by regulating Sox2 expression, thereby enabling cell shape changes crucial for neural tube morphogenesis (Fig. 6).

This study underscores the critical role of neural activity and signaling during neural tube formation. Thus, the neural plate and the formation of the neural tube are vulnerable to exposure of environmental factors that can disrupt this neural signaling. Our study provides a foundational understanding of the mechanisms involved in neural tube morphogenesis, which can serve as a mechanistic basis to further investigate how neuroactive drugs interfere with this process, thereby advocating for the cautious use of such therapeutics during pregnancy.

## Materials and Methods

### Animals

All experimental procedures and research design utilized in this study complied with ethical regulations. The Institutional Animal Care and Use Committee approved the animal protocol #22264 implemented in this study. IACUC follows the guidelines established by the Animal Welfare Act and the Public Health Service Policy on Humane Care and Use of Laboratory Animals.

### *Xenopus laevis* animal handling and *in vitro* fertilization

Mature oocytes were collected in a dish from a previously hCG injected female frog and incubated with a small piece of minced testis. This is considered time 0 of fertilization. Fertilized oocytes were kept in 10% MMR saline, containing (in mM): 10 NaCl, 0.2 KCl, 0.1 MgSO_4_, 0.5 Hepes, 5 EDTA and 0.2 CaCl_2_. Dejellying of embryos was done by briefly swirling fertilized eggs in 2% cysteine solution, pH 8. Developmental stages were recorded according to Nieuwkoop and Faber ^52^. Animals were handled according to the IACUC guidelines using humane procedures to prevent animal suffering.

## Gene expression assays

Neural plates from 9 mid neural plate stage (stage 16) *Xenopus laevis* embryos were isolated, resuspended in Trizol reagent (Invitrogen, cat. # 15596026) and stored at -80°C. RNA was extracted with kit according to manufacturer’s instructions (RNeasy Mini Kit, Qiagen, cat. # 74104), gDNA was eliminated (RapidOut DNA Removal Kit, Thermo Scientific, cat. # 00859896) and cDNA was made (High-Capacity cDNA Reverse Transcription Kit, Applied Biosystems, cat. # 00890068) with standard protocols. Using this cDNA as template, RT-PCR was performed with the following primers: vesicular glutamate transporter 1 (*vglut1*): forward, GCAACTTGGGTGTAGCCATT, reverse, TGCCCATTTACTCCAGATCC; excitatory amino acid transporter 5 (*eaat5*): forward, GTGGGATGTCTGCTTGGATT, reverse, ATGTGGCTTCCACAAGGTTC; syntaxin 1A (*stx1a*): forward, ATGAAGGATCGGACCAGGGA, reverse, TGTGGCGTTGTATTCGGACA; vesicle associated membrane protein 1 (*vamp1*): forward, GCCACAGGTGATCCTGGAAA, reverse, AGGAGACGCTCCACACAATG; synaptosome associated protein 25 (*snap25*) forward, AAGGCTTGGGGCAATAACCA, reverse, AACCACTGCCCAGCATCTTT, designed using Primer-BLAST (www.ncbi.nlm.nih.gov). All sequences are written from 5’ to 3’.

## VGluT1 knockdown

Two-cell-stage embryos were unilaterally or bilaterally injected with 2-4 pmol of translation blocking morpholino (MO) VGluT1-MO1 TCCTAAACTCCATTGTGATCCTCCT (VGluT1-KD1/KD) or with 8 pmol splicing blocking MO (VGluT1-KD2) VGluT1-MO2 ATGTTTTTTCCTTACCTCGATAACA per blastomere (Gene Tools, Inc). Controls were sibling embryos injected with standard control MO, CCTCTTACCTCAGTTACAATTTATA (Control). Morpholinos were injected along with dextran-Alexa-Fluor conjugates or with GFP or mCherry mRNA to assure permanency of MO reporter after PFA or TCA fixation. All sequences are written 5’ to 3’. Rescue experiments were implemented by injecting 250 pg of VGluT1-MO-resistant *Xenopus laevis vglut1*- mRNA (125 pg/blastomere) along with VGluT1-MO. *X. laevis* VGluT1 mRNA was modified by substitution of 5’ UTR with a fragment of the Kozak consensus sequence gcc acc (Kozak, 1987). A number of wobble mutations were introduced in the VGluT1 coding region to prevent VGluT1-MO binding to modified *vglut1* mRNA. mRNA was synthesized as previously described ^30,53–55^. Assessment of VGluT1-KD efficiency was performed by Western blot assays.

## Western blot assays

To determine VGluT1 and Sox2 protein levels, neural tube, stage-22 whole embryos were homogenized in extraction buffer containing 1% Triton X-100, 150 mM NaCl, 25 mM Hepes pH 7.4, 2 mM EDTA and protease inhibitors cocktail (Thermo Fisher Scientific). Samples were centrifuged at 16,100 *g* for 10 min and the pellet discarded. Supernatant was then boiled with Laemmli buffer and run on a 10% SDS-PAGE followed by protein transfer to PVDF membrane. PVDF membranes were probed with monoclonal anti-*Xenopus*- VGluT1 1:500 (Genescript, costume-made) 5% milk, Sox2 1:500 (catalog# AF2018 R&D Systems) in 5% BSA followed by incubation with HRP-conjugated secondary antibodies 1:10,000 (Jackson ImmunoResearch) and visualized by Western Lightning Plus-ECL, Enhanced Chemiluminescence Substrate (catalog# NEL103E001, PerkinElmer). PVDF membranes were stripped in 0.2 M glycine HCl buffer, pH 2.5, 0.05% Tween for 20 min and reprobed with GAPDH 1:50,000 in 5% Milk (catalog# 60004-1-1g Proteintech), as loading control, followed by incubation with HRP-conjugated secondary antibodies 1:20,000 (Jackson ImmunoResearch) and visualized by Western Lightning ECL (Millipore/Sigma, cat. # GERPN2106).

## Whole-mount immunostaining

Embryos at stage 13 through 18 were fixed in 4% PFA for 2 h at 23°C or overnight at 4°C and the dorsal half containing the neural plate was isolated and bleached in 1:2 Dent’s fixative/H_2_O_2_ overnight at 23°C. Samples were washed, permeabilized in 1% Triton-X100, and incubated overnight at 4°C with primary antibodies, followed by staining with fluorescent secondary antibodies at 23°C for 2 h, and finally clearing overnight in benzyl benzoate. Z-stack confocal images of embryos or neural tissue (100 μm-thick) were taken with a confocal microscope (Nikon C1 or C2), 10X or 20X objective, through approximately 30-100 steps (3-10 μm) longitudinally through a dorsoventral direction. Primary antibodies used for whole mount immunostaining were: VGluT1 1:200 (catalog # AB5905, EMD-Millipore), Sox2 1:200 (catalog # AF2018, R&D Systems), pERK1/2 1:200 (catalog # 4377S, Cell Signaling Technology), PCNA 1:1,000 (catalog # 2586S, Cell Signaling Technology), PHH3 1:500 (catalog #06-570, EMD Millipore), and GFP 1:500 (catalog # ab13970 Abcam).

Quantitative assessment of the number of cells immunopositive for VGluT1, Sox2, PCNA and PHH3 was performed by using the Imaris ‘Spot’ function to detect nuclei objects filtered by object size, fluorescence intensity, and the built-in quality threshold.

## Electron Microscopy

### Transmitted EM

Embryos were fixed in 3% glutaraldehyde in 100 mM HEPES (pH 7.4) overnight at 4°C and washed several times in 100 mM HEPES and processed for routine TEM. Specimens were postfixed in 1% OsO4 in PBS overnight on ice and washed in PBS and water. Fixed embryos were embedded in epoxy blocks. Ultrathin sections were stained with uranyl acetate and lead citrate prior to viewing on a transmission electron microscope (Philips CM120 Biotwin Lens, FEI Company) using a Gatan MegaScan digital camera (model 794/20, 2K × 2K, Gatan).

### Immuno-TEM

Embryos were fixed in 4% paraformaldehyde in 0.1 M sodium phosphate buffer at 4°C overnight. Ultrathin sections were incubated with primary and secondary antibodies overnight at 4°C and for 2 h at 23°C, respectively. Primary antibody was VGluT1 1:100-1:10,000 (catalog # AB5905, EMD-Millipore). Secondary antibody was goat-anti-guinea pig IgG, EM grade, 10 nm 1:40 (catalog # 25329, Electron Microscopy Sciences). Immunolabeled sections were imaged on the same electron microscope as indicated before.

## Measurement of glutamate release

### iGluSnFR

Two-cell stage wild type and unilaterally injected with 2 pmol VGluT1-MO1 or Control-MO+Alexa 647- dextran embryos were bilaterally injected with 8 ng iGluSnFR mRNA per embryo [iGluSnFR, a modified (pcDNA-spacer-123D5, Lin Tian’s Lab) derived from construct gift from Loren Looger, Addgene plasmid #41732, ^56^] that was previously subcloned into the pCS2^+^ vector. iGluSnFR-expressing stage 13-16 embryos were confocally time-lapse imaged at 1 Hz acquisition rate. Images were taken at different neural plate stages using Nikon confocal microscope in response to 2-5 μM ionomycin or DMSO for 5 min followed by 5 mM glutamate. Membrane-mCherry expressing embryos were used as controls. Fluorescence intensity was measured using NIS Elements software.

### Fluorometric assay

Neural plate stage (stage 15-15.5) wild type and morpholino-injected embryos were incubated with 100 nM tetanus toxin or DMSO for 1 h followed by 1 h incubation with 2 μM ionomycin in 200 μl volume. Bathing solution was collected and released glutamate measured with a Fluorometric Glutamate Assay Kit (STA-674 Cell-biolabs).

### *In vivo* Ca^2+^ imaging

DNA encoding the Ca^2+^ sensor GCaMP6s (pGP-CMV-GCaMP6s, a gift from Douglas Kim, HHMI Janelia Research Campus, Ashburn, Virginia; plasmid #40753, Addgene; (Chen et al., 2013)) was subcloned into the pCS2^+^ vector using BglII and NotI restriction sites. The BglII restriction site was included in pCS2^+^ with the following primers: forward, 5’ -TCACTAAAGGGAACAAAAGATCTGGGTACCGGGCCCAA-3’; reverse, 5’ - TTGGGCCCGGTACCCAGATCTTTTGTTCCCTTTAGTGA-3’. For all experiments in this study, mRNA was transcribed from the indicated plasmids using mMessage mMachine kits (Ambion). GCaMP6s mRNA was injected in two-cell-stage embryos (1 ng mRNA/embryo). Neural plate stage embryos [14 –19 h postfertilization (hpf)] were imaged under a confocal microscope at an acquisition rate of 0.05-0.1 Hz for 5 min to 2 h. Detection of Ca^2+^ transients was thresholded by a peak change in fluorescence of at least 2 times the noise, as in previous studies ^53,54,57^. The number of Ca^2+^ transients in unilaterally VGluT1-KD1 or Control embryos were compared, and significance was assessed by paired t test.

### Imaging of Erk1/2 activity

piRFP670 (a gift from Vladislav Verkhusha; Addgene plasmid # 45457; ^58^) was subcloned into pCS2^+^ using KpnI and NotI (New England Biolabs; NEB). The ERK-sensitive region of ERKKTRClover from pENTRA- ERKKTRClover (a gift from Markus Covert; Addgene plasmid # 59138; ^38^) was removed and amplified by PCR using the following primers: forward: CAAAGGTACCGGCAACATGGCAAAGGGCCGAAAGCC, reverse: CACCATACCGGTGAGGATGGGAATTG. These primers add KpnI and AgeI restriction sites, keep the iRFP in frame with the KTR, and optimize the Kozak sequence of the new construct for *Xenopus laevis* expression ^59,60^. The PCR product and piRFP670-pCS2^+^ were both digested with KpnI and AgeI (NEB), gel purified, and ligated. The new plasmid was confirmed by restriction digests and RNA was made from the T3 promoter using Ambion’s mMessage mMachine kit. The sensitivity of the new construct was confirmed in culture as described below using phorbol 12-myristate 13-acetate (PMA; agonist) or PD0325901 (antagonist).

Embryos were injected at the 2-cell-stage with a mixture of mRNA encoding GCaMP6s (1.5 ng), ERK-KTR- iRFP670 (1.5 ng), and H2B-mRFP1 (200 pg; mRNA made from the SP6 promoter of pCS2^+^-H2B-mRFP1, a gift from Sean Megason; Addgene plasmid # 53745), or 2 pmol VGluT1-MO or Control-MO (unilateral; along with tracer), 1.5 ng ERK-KTR-iRFP670 (bilateral), and 200 pg H2B-mRFP1 (bilateral). They were grown at 18-21°C until early neural plate stages when they were confocally imaged in the 3 channels.

For simultaneous Ca^2+^ and Erk1/2 activity analysis, nuclei were tracked by H2B signal in NIS Elements software (Nikon) and the intensity of all three channels in this H2B-RFP-fluorescently-labeled area (nuclear) and surrounding area (cytosolic) over time were measured. Cells were categorized into Ca^2+^- active or Ca^2+^-silent based on occurrence of Ca^2+^ transients or not during the recording, respectively. At least 5 cells of each category were analyzed per embryo (total: 70 cells). Ca^2+^ active cells were first selected and Ca^2+^-silent cells were sampled to match anteroposterior and mediolateral contralateral localization to those active in each embryo. Average of cytosolic/nuclear ERK-KTR-iRFP670 was calculated per group per embryo and comparisons were performed with paired t-test.

For assessing Erk1/2 activity dependence on VGluT1 expression we imaged embryos at mid-neural plate stages (stage 15-16) and measured ERK-KTR-iRFP670 cytosolic/nuclear signal in cells of VGluT1- MO/Control-MO injected and wild type halves of neural plate. Cells were sampled to match anteroposterior and mediolateral localization across groups. Average of cytosolic/nuclear ERK-KTR- iRFP670 of at least 5 cells in each half of the neural plate from at least 4 embryos (total neural plate cells at least 40) was calculated per wild type and injected neural plate per embryo and comparisons were performed with paired t-test.

### Immunohistochemistry of neural tissue thin sections

Neural plate stage embryos were fixed at 23°C with 4% PFA for 10 min, and processed for immunostaining as previously described ^55^. Briefly, samples were paraffin-embedded and 12 μm-thick transverse sections of the neural plate were incubated with primary and secondary antibodies overnight at 4°C and for 2 h at 23°C, respectively. Primary antibody used was: Myc-tag 1:2,000 (catalog # 2276, Cell Signaling). Antigen retrieval was performed by boiling samples in 0.05% citraconic anhydride, pH 7.4 for 10 min in water bath^61^. Samples were permeabilized with PBST (0.5% Triton) for 1 h at 23°C, blocked with 5% BSA in PBST (0.1% Triton) for 30 min using SNAP i.d. 2.0 System for immunohistochemistry (Millipore). Samples were imaged with a confocal microscope (Nikon A1), 40X objective through approximately 15 1-μm steps.

### VGluT1 knockout

Synthetic guidance RNA (sgRNA) targeting *vglut1* was designed using the CRISPRscan website ^62^, and inDelphi model ^63^, which provide in silico predictions for mutational outcomes. The VGluT1 sgRNA (guidance sequence: AGCCTGCTACGCTCCAGAGG) was synthesized using the EnGen sgRNA synthesis kit (New England Biolabs). The sgRNA was complexed with Cas9 protein (PNA Bio, #CP02) at 300 mM KCl to form ribonucleoprotein complexes and injected into embryos. The CrispantCal web tool ^64^ was used to calculate volumes corresponding to an optimal one-to-one molecular ratio of gRNA to Cas9 in a CRISPR- Cas9 injection mix. To quantify the editing efficiency, genomic DNA was extracted from 5 edited embryos at early neural plate stages (stage 14) using DNeasy Blood & Tissue Kit (Qiagen). The edited locus of *vglut1.s* was amplified from genomic DNA using primers, forward, 5’-TTTGCGCTTGACCCAGGTAT-3’; reverse, 5’-ACGGGCGACAATTTTATGCG-3’, specific to the CRISPR/Cas9-targeted site for Sanger sequencing. The sequencing results were used for in silico analysis of the INDELs generated by the CRISPR/Cas9- mediated editing using Inference of CRISPR Edit analysis software (Synthego).

### Assessment of Neural Tube Defect phenotype

VGluT1-KD, VGluT1-KO, Control and vglut1-mRNA injected embryos were imaged with a stereoscope when control embryos reached early neural tube stages (stage 20). Observed phenotypes were categorized in “Closed” or open (“NTDs”) neural tube.

### Statistics & Reproducibility

Rigorous research design and analysis was implemented by running all the controls necessary alongside experimental samples. Data analysis was performed blindly by the experimenter or through unbiassed automation. Number of samples for each experiment was determined by power analysis of pilot experiments and replicated at least 3 times with at least 5 samples. No data were excluded from the analysis. Statistical analysis of the data was done with Prism software (Graphpad, Inc.). Normality test was performed in each set of data and then parametric (normally distributed) or non-parametric statistical analysis was chosen. Paired tests were implemented in unilaterally manipulated embryos (comparison of non-injected and microinjected halves of neural tissue), or when Ca^2+^ activity or released glutamate (iGluSnFR) was compared before and after addition of an agent in the same sample. Differences were considered statistically different when p<0.05. Exact p values are included in the Source Data File for each data set. Type of statistical test used is indicated in the figure legends.

## Acknowledgments

This work was supported by: National Science Foundation grant 1754340 (LNB), National Institutes of Health grants R01NS105886 and R01NS113859 (LNB), Shriners Hospital for Children grant 85111 (LNB) and grant 84303 (RG)

## Author contributions

Conceptualization: RG, LNB Methodology: RG, PAC, SS, JL, GOM Investigation: RG, PAC, JL Funding acquisition: RG, LNB Project administration: LNB Supervision: LNB, LT Writing – original draft: RG, LNB Writing – review & editing: RG, JL, SS, LNB

---

**Extended Data Figure 1.**
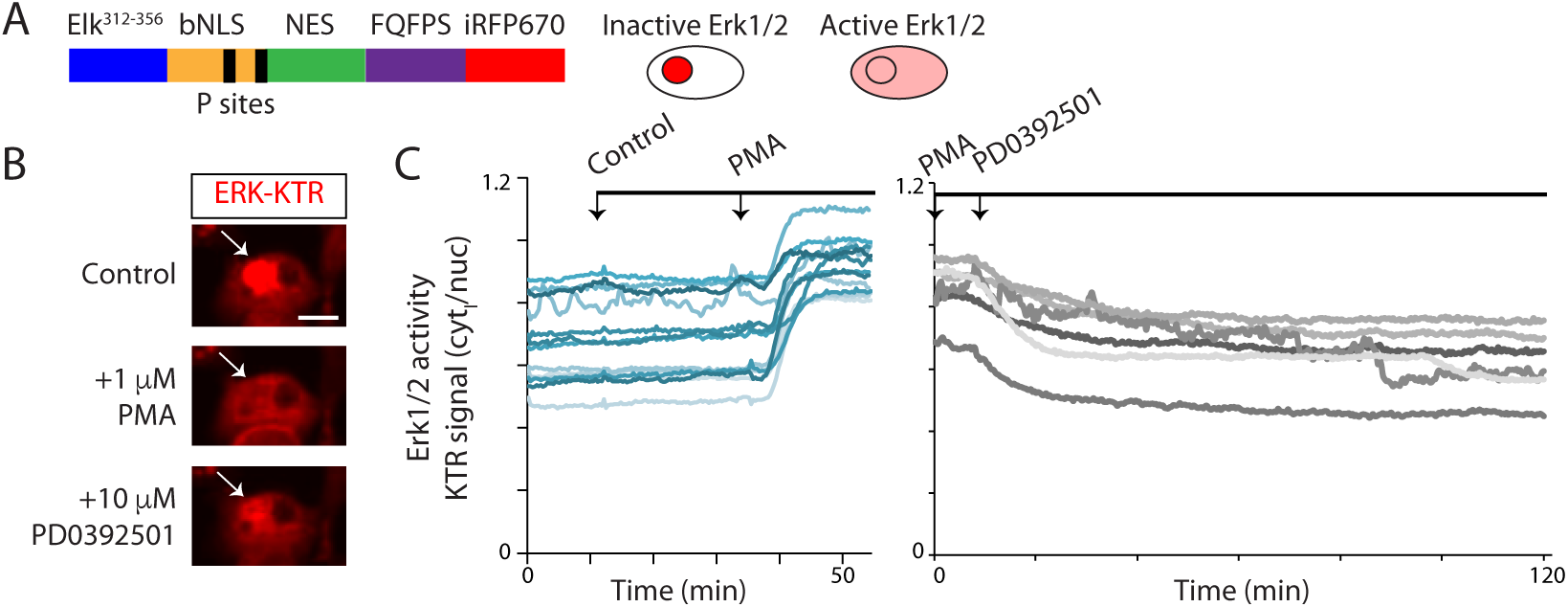
**ERK-KTR reports Erk1/2 activity in *Xenopus laevis* embryonic cells**. Muscle and spinal cord were dissected from neural tube stage embryos expressing the Erk1/2-kinase translocation reporter (ERK- KTR), a reporter of Erk1/2 activity that is exported out of the nucleus when Erk1/2 is active. Cells were dissociated and plated *in vitro*. One day after plating, cells were time-lapse imaged with a rate of 0.05 Hz. **A.** Schematic shows ERK-KTR construct design. Black bars represent Erk1/2 specific phosphorylation sites. Cell drawings depict subcellular localization of ERK-KTR reporter in cells with lower (inactive) and higher (active) levels of Erk1/2 activity. **B-C.** To test the specificity of the reporter, cells were incubated in vehicle (0.1% DMSO; Control), 1 µM phorbol-12-myristate-13-acetate (PMA; an activator of MAP kinases including Mek1/2), or 10 µM PD0392501 (Mek1/2 inhibitor). **B.** Images show a representative example of a developing neuron before treatment, after PMA, and after PD0392501 additions. Scale bar, 10 μm. Arrow points to the nucleus. **C.** Graphs show examples of the change in the ratio of cytosolic (cyt) to nuclear (nuc) fluorescence in individual cells in representative experiments. Arrows indicate the time of addition of indicated agent. N ≥4 cell cultures, n ≥24 cells.

**Extended Data Figure 2.**
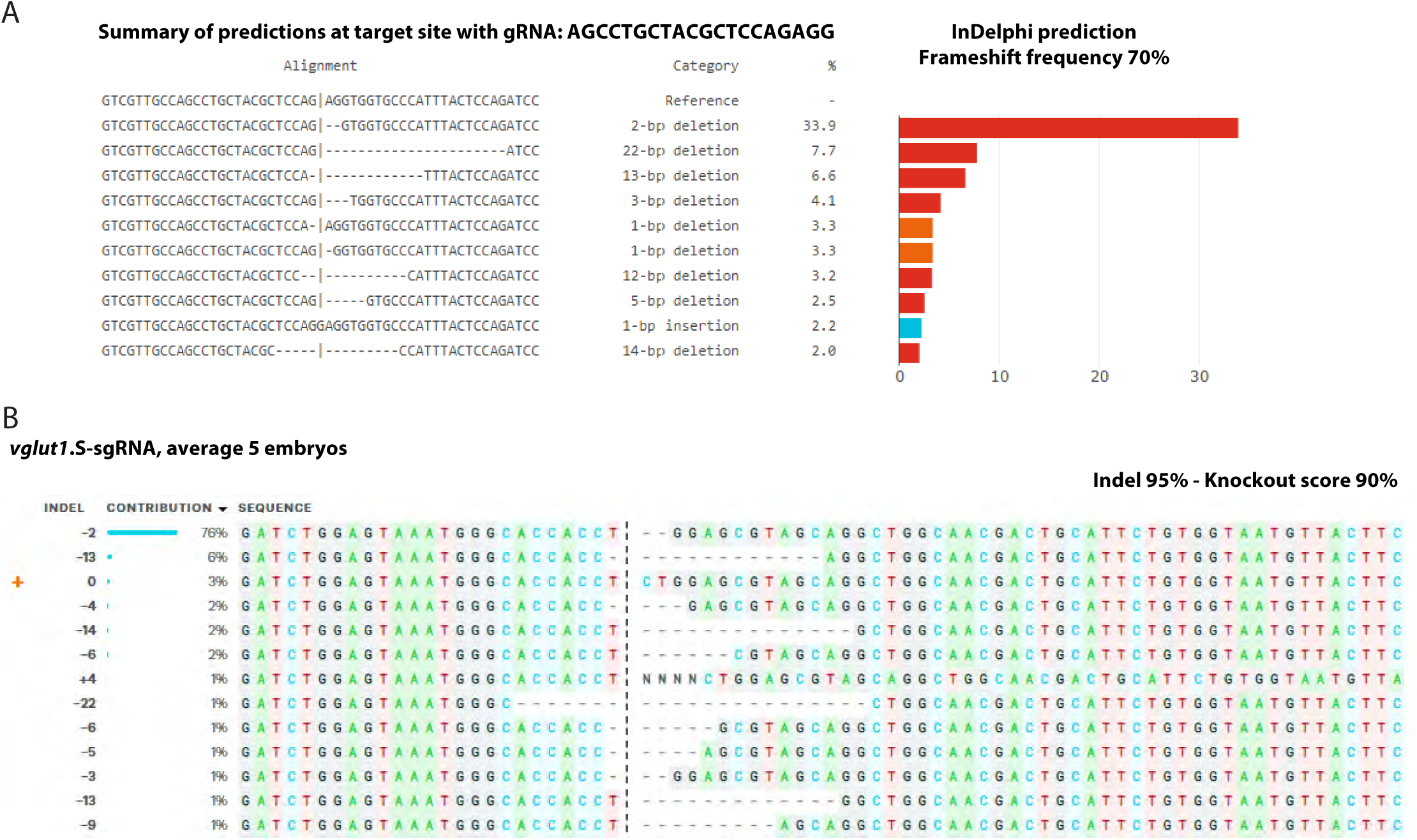
Prediction and experimental outcomes of CRISPR/Cas9 gene editing against *vglut1* in *Xenopus laevis* embryos. A. InDelphi prediction of *vglut1*.S CRISPR/Cas9-mediated editing outcome. B. Two-cell stage embryos were unilaterally injected with 1 ng Cas9 protein and 0.5 ng sgRNA against *vglut1*.S and processed for sequencing and ICE knockout analysis at stage 22. Experimental assessment of CRISPR/Cas9-mediated *vglut1.S* gene deletion by ICE knockout analysis shows 95% Indel and 90% knockout score. The Cas9 cleavage site is indicated in vertical dashed line. Wild-type sequence is marked by + (left, 0).

