## Extended Data Figs 1-2 for "Vesicular glutamate release is necessary for neural tube formation"

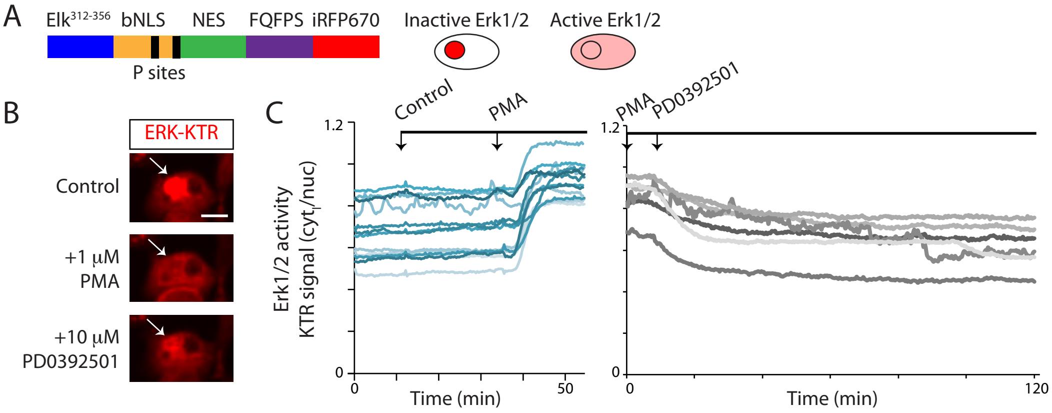

A
Elk312-356
bNLS
NES
FQFPS
iRFP670
P sites
Inactive Erk1/2
Active Erk1/2
B
ERK-KTR
Control
+1 μM PMA
+10 μM PD0392501
C
Control
PMA
PMA
PD0392501
Erk1/2 activity
KTR signal (cyt/nuc)
1.2
0
0
50
Time (min)
1.2
0
0
120
Time (min)

A

**Summary of predictions at target site with gRNA: AGCCTGCTACGCTCCAGAGG**

| Alignment | Category | % |
|---|---|---|
| GTCGTTGCCAGCCTGCTACGCTCCAG\|AGGTGGTGCCCATTTACTCCAGATCC | Reference | - |
| GTCGTTGCCAGCCTGCTACGCTCCAG\|--GTGGTGCCCATTTACTCCAGATCC | 2-bp deletion | 33.9 |
| GTCGTTGCCAGCCTGCTACGCTCCAG\|----------------------ATCC | 22-bp deletion | 7.7 |
| GTCGTTGCCAGCCTGCTACGCTCCA-\|------------TTTACTCCAGATCC | 13-bp deletion | 6.6 |
| GTCGTTGCCAGCCTGCTACGCTCCAG\|---TGGTGCCCATTTACTCCAGATCC | 3-bp deletion | 4.1 |
| GTCGTTGCCAGCCTGCTACGCTCCA-\|AGGTGGTGCCCATTTACTCCAGATCC | 1-bp deletion | 3.3 |
| GTCGTTGCCAGCCTGCTACGCTCCAG\|-GGTGGTGCCCATTTACTCCAGATCC | 1-bp deletion | 3.3 |
| GTCGTTGCCAGCCTGCTACGCTCC--\|----------CATTTACTCCAGATCC | 12-bp deletion | 3.2 |
| GTCGTTGCCAGCCTGCTACGCTCCAG\|-----GTGCCCATTTACTCCAGATCC | 5-bp deletion | 2.5 |
| GTCGTTGCCAGCCTGCTACGCTCCAGGAGGTGGTGCCCATTTACTCCAGATCC | 1-bp insertion | 2.2 |
| GTCGTTGCCAGCCTGCTACGC-----\|---------CCATTTACTCCAGATCC | 14-bp deletion | 2.0 |

**InDelphi prediction**
**Frameshift frequency 70%**

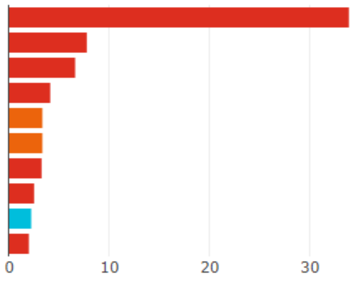


B

***vglut1*.S-sgRNA, average 5 embryos**

**Indel 95% - Knockout score 90%**

| | INDEL | CONTRIBUTION | SEQUENCE |
|---|---|---|---|
| | -2 | 76% | GATCTGGAGTAAATGGGCACCACCT ¦ --GGAGCGTAGCAGGCTGGCAACGACTGCATTCTGTGGTAATGTTACTTC |
|  | -13 | 6% | GATCTGGAGTAAATGGGCACCACC- ¦ ------------AGGCTGGCAACGACTGCATTCTGTGGTAATGTTACTTC |
| + | 0 | 3% | GATCTGGAGTAAATGGGCACCACCT ¦ CTGGAGCGTAGCAGGCTGGCAACGACTGCATTCTGTGGTAATGTTACTTC |
| | -4 | 2% | GATCTGGAGTAAATGGGCACCACC- ¦ ---GAGCGTAGCAGGCTGGCAACGACTGCATTCTGTGGTAATGTTACTTC |
|  | -14 | 2% | GATCTGGAGTAAATGGGCACCACCT ¦ --------------GCTGGCAACGACTGCATTCTGTGGTAATGTTACTTC |
|  | -6 | 2% | GATCTGGAGTAAATGGGCACCACCT ¦ ------CGTAGCAGGCTGGCAACGACTGCATTCTGTGGTAATGTTACTTC |
| | +4 | 1% | GATCTGGAGTAAATGGGCACCACCT ¦ NNNNCTGGAGCGTAGCAGGCTGGCAACGACTGCATTCTGTGGTAATGTTA |
|  | -22 | 1% | GATCTGGAGTAAATGGGC------- ¦ ---------------CTGGCAACGACTGCATTCTGTGGTAATGTTACTTC |
|  | -6 | 1% | GATCTGGAGTAAATGGGCACCACC- ¦ -----GCGTAGCAGGCTGGCAACGACTGCATTCTGTGGTAATGTTACTTC |
|  | -5 | 1% | GATCTGGAGTAAATGGGCACCACC- ¦ ----AGCGTAGCAGGCTGGCAACGACTGCATTCTGTGGTAATGTTACTTC |
| | -3 | 1% | GATCTGGAGTAAATGGGCACCACC- ¦ --GGAGCGTAGCAGGCTGGCAACGACTGCATTCTGTGGTAATGTTACTTC |
|  | -13 | 1% | GATCTGGAGTAAATGGGCACCACCT ¦ -------------GGCTGGCAACGACTGCATTCTGTGGTAATGTTACTTC |
|  | -9 | 1% | GATCTGGAGTAAATGGGCACCACCT ¦ ---------AGCAGGCTGGCAACGACTGCATTCTGTGGTAATGTTACTTC |
